# DeepRod: A human-in-the-loop system for automatic rodent behavior analysis

**DOI:** 10.1101/2024.01.04.572506

**Authors:** A. Loy, M. Garafolj, H. Schauerte, H. Behnke, C. Charnier, P. Schwarz, G. Rast, T. Wollmann

## Abstract

We present a human-in-the-loop system for efficient rodent behavior analysis in drug development. Addressing the time-consuming and labor-intensive nature of manual behavior categorization, this UX-optimized platform integrates AI for complex behavior prediction and active learning to identify rare events. The proposed solution leverages a cloud-native data processing pipeline, AI-based novelty behavior recognition and multi-class classification, demonstrating significant improvements in behavior labeling and discovery.

## Introduction

In drug discovery and development, systematic assessment of drug safety using highly regulated preclinical studies prior to first-in-human clinical trials are mandatory [1] to ensure safety for volunteers and patients. These assessments allow detailed views on risk, benefit and the therapeutic index of potential future therapeutics. The evaluations include, among others, standardized functional behavioral studies in rodents [2, 3, 4]. During the research phase, automated video-based systems (e.g., PhenoTyper) to assess continuous quantitative and qualitative motor behavior during the active phase of the rodents. Infrared video cameras located in the top unit of each observation arena populated with one rodent per arena record 14 h of video material per animal. During each study, groups of animals exposed to an active ingredient at various doses or receiving a placebo are recorded from the top, generating large video datasets. Events of interest can be very rare and require in depth analysis of the footage. Manual analysis is not feasible in an appropriate time and with acceptable effort.

Automated analysis of distance moved and animal velocity (e.g. Ethovision XT) provides very sensitive measures for central nervous effects and general tolerability. However, these features are not discriminative enough to detect complex events.

In this work, we propose an UX-optimized platform for behavior labeling and analysis integrated into the workflow, AI-based complex behavior prediction, active learning to find rare events and propose candidates for new behavioral categories.

## Methods

We propose a novel system for analyzing rodent behavior at scale, which combines a user-centered interface with AI-based behavior prediction, novel behavior recognition and active learning. The system should support rodent behavioral analysis by automatizing the behavior classification of rodents. To achieve this, the system needs to enable users in annotating the rodent behavior in video snippets to gather training data, support users in detecting novel, previously unseen behaviors and finally to automatically classify the behavior of rodents.

### System overview

Each routine study performed to profile a research substance includes 35 million frames from typically 28 individual 14h long video clips with 25 fps, recorded by a PhenoTyper camera [5, 6]. To tackle this amount of data, an efficient, parallelized, and cloud-native data processing pipeline processes these raw video files [7]. First, the pipeline re-encodes the input video for efficiency and storage and registers the video’s metadata into the system. Second, visual information is extracted using a deep learning approach [8] and meaningful features are further generated using that information. Lastly, a classifier predicts the rodent behavior for each frame based on those features.

### Behavior recognition

The core component of our system is the behavior recognition component. This component uses a two-staged machine learning pipeline to classify the behavior of rodents. The first stage extracts visual information from the video stream by localizing nine anatomical landmarks (“keypoints” in the following) of the rodent. Similar to MARS [9], keypoints correspond to the nose, ears, body center, hips, tail base, tail center and tail end. Our keypoint extraction method is based on DeepLabCut [8].

The second stage is a classifier that uses features based on the keypoints. We identified that features capturing the position, pose, and movement are discriminative for characterizing the rodent’s behavior. Per each feature category, we engineered a range of features based on the keypoints capturing the rodent’s position, pose and movement (Figure 1). Some features are aggregated within sliding windows of various sizes. We frame the behavior detection problem as a multi-class classification problem, containing all known behaviors and an extra class representing any unknown behavior, which is explicitly labeled to not conform to any of the known behavior. Our method is leveraging XGBoost [12], which is recognized as a good classifier under skewed data and noise [13]. The tree-based model also enables interpretability like computing feature importance, which is favorable in life sciences [14]. The system trains new models automatically based on user request. Users receive a report after training that offers an intuitive overview of model improvements.

**Figure 1.1.**
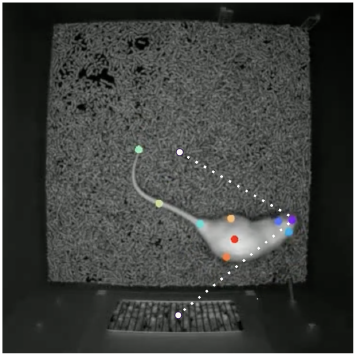
Features based on the distances between keypoints are relevant to detect interactions.

**Figure 1.2.**
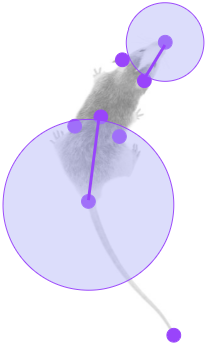
Features based on [10] use the relative keypoint positions are relevant to identify typical postures.

**Figure 1.3.**
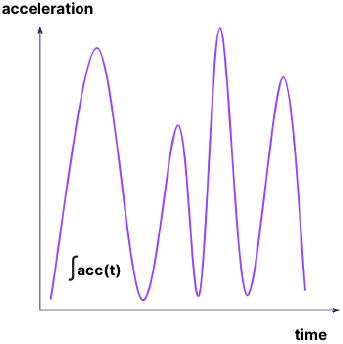
Features based on [11] use the keypoint movements to identify temporal patterns like directed or undirected motion.

### Detection and labeling of rare behavior

To enable automatic rodent behavior classification, collection of annotated data is necessary to train the machine-learning-based classifier. As there are thousands of hours of video material that can be used to create the annotated training data set, the choice of which video sections to annotate is not trivial and is subjected to time constraints of the labeling force.

For efficient use of human labeling resources, the system implements a labeling assistant shown in Figure 2. The labeling assistant leverages an active learning method based on Meal [15] to suggest areas to label across the whole video material. The active learning methodology can be formulated as an SQL query, which selects a fixed amount *k* of currently unlabeled model predictions from the database. We refer to the result of this query for a given video as the “labeling queue”. The active learning query consists of multiple subqueries, which each have a capped contribution to the overall labeling queue. The subqueries select samples with the following properties: Label is likely to belong to an underrepresented class in the annotated data set, has high prediction uncertainty, has high novelty score, is selected randomly.

**Figure 2.**
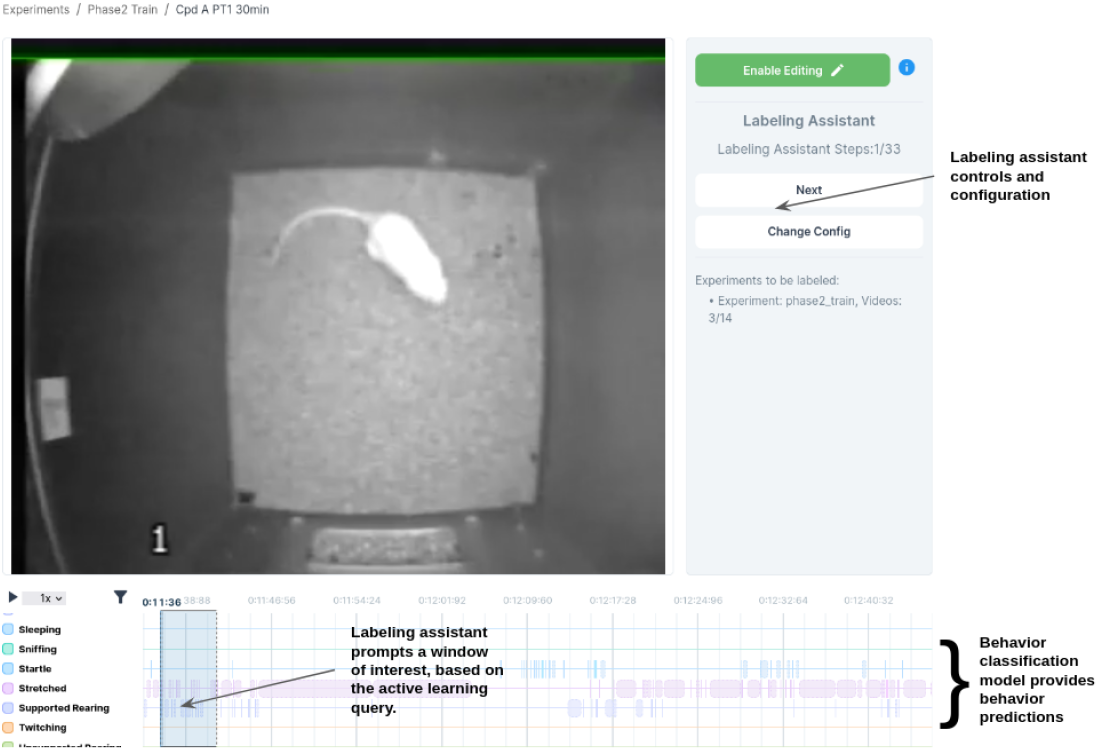
Main view of annotation view with the labeling assistant. Labelers are presented with the video footage and the timeline of model-predicted behaviors as well as annotations that are already set. Annotations can be created with a single click to allow for an efficient process.

Each item in the labeling queue corresponds to a window of interest, which users are expected to annotate. The order of the queue is randomized to prevent bias of which sections in the video get labeled. Once the queue is exhausted, labelers are directed to a new video with a fresh queue and therefore iteratively cycle through all videos. This setup prevents annotations concentrated in only a small subset of videos and ensures that each video acquires labels for the top *k* most relevant sections.

### Novel behavior recognition

The rodents in the experiments might demonstrate unusual or novel behaviors due to the effect of the compounds that they are exposed to. Therefore, the system needs to support users in observing unusual behavior to enable them to possibly categorize it as a new behavior class. We refer to this problem as **“**novel behavior recognition” and formulate it as an outlier detection problem. Each frame gets embedded into a low-dimensional feature space using principal component analysis (PCA) to reduce the dimensionality and the correlation between features. Then, mean and variance across all labeled samples for each class (i.e. each defined and annotated behavior type) are computed. With that, we define the *novelty score* of a frame as the *Mahalanobi’s distance* [16] to the closest known distribution.

## Results

The system was evaluated in a pilot with three distinct expert annotators who created 13.862 new annotations. In this context, one annotation refers to an identified behavior type with a start and end frame. These annotations are distributed across 226 individual rodents from 16 distinct experiments.

### Active learning results

With the help of the active learning component highlighting areas of high interest, the experts identified and added several new behavior types. Figure 4 demonstrates the benefit of using the system to extend the training dataset.

The novelty behavior detection model was evaluated through a leave-one-out assessment due to the absence of explicit labels for training and evaluation. This approach involves iteratively treating each known rodent behavior as novel, allowing us to gauge the proficiency of the method in identifying these established behaviors as potentially new instances. Further, it gives insights into the ability to expedite the discovery of behaviors by ranking them higher in the novelty queue. This is crucial for labelers using the labeling assistant algorithm, as it aids in identifying behaviors promptly rather than randomly later in the process. We observed a significant improvement in 6 out of 9 behavior types. Some of the behavior types would not have been prioritized by the novelty ranking, largely due to their intermediate position in the reduced feature space (Figure 3). Thus, we extended the active learning component to be composed of multiple strategies additionally to novelty ranking.

**Figure 3.**
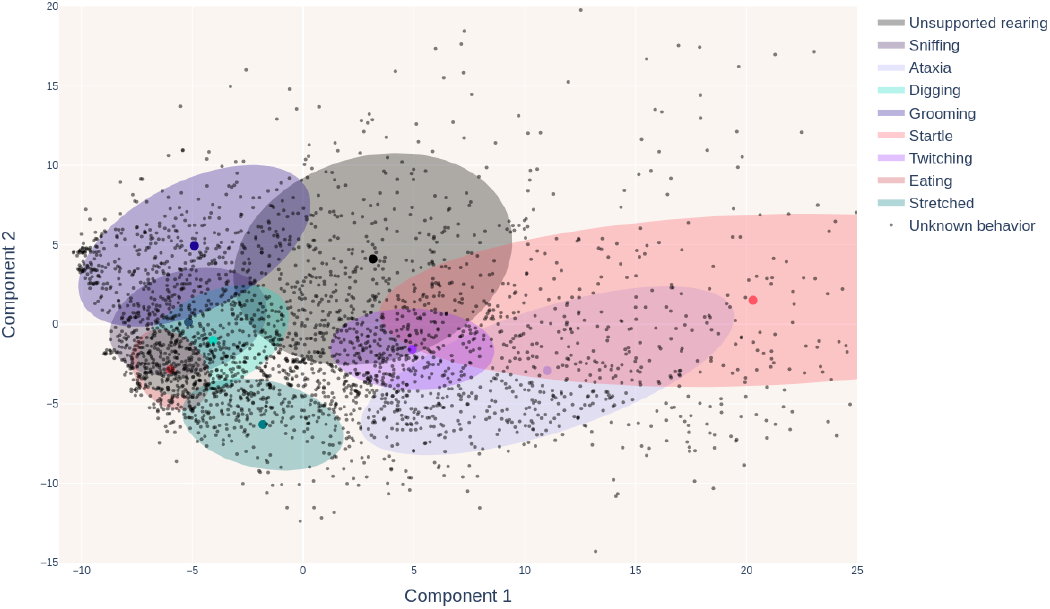
Visualization of the lower dimension feature embedding space used for novelty scoring of a sample. We visualize the center and variance of the estimated class distributions. Samples are scored according to their distance to the distributions of labeled samples. Note that the figure contains a subset of randomly sampled 3000 unlabelled behavior points.

**Figure 4.**
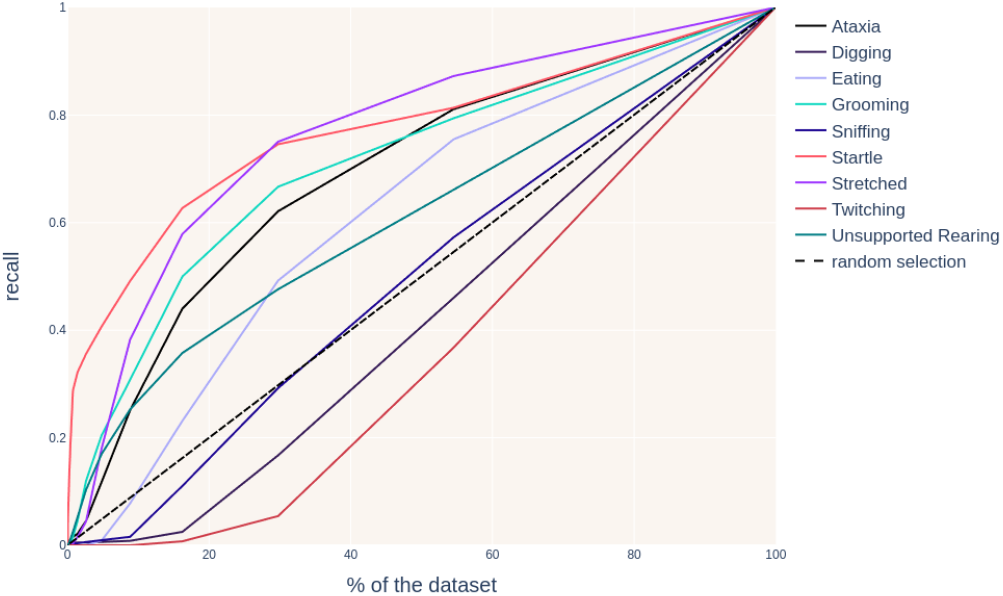
Recall of our system for rare behavior types with different subsets of the dataset. The percentage of coverage of a novel behavior when annotating a certain percentage of the overall dataset is shown.

### Classification results

As the amount of behavior types was heavily extended during the project, a direct comparison of model results at the beginning and the end of the project is not feasible. However, a strong improvement can be seen in behavior types for which little training data was available at the start of the project due to the rare occurrence of these behaviors as shown in Table 1. With the data collection using our system and active learning, the amount of labels for some of these behavior types could be increased substantially. This enabled the training of a model, which can detect these behaviors more reliably. Examples for this are “Grooming”, where the true positive (TP) rate improved from 6% to 73%, and “Twitching” with an improvement from 3% to 29%. Moreover, the final model is able to detect a larger number of distinct behavior types.

**Table 1.** Comparison of the performance between the initial model and the final model, as well as the difference in data set size. Each row refers to a class of behavior type that the models aim to detect. TP refers to the true positive rate of the predictions, evaluated on a per-frame basis. Label Data Increased lists the factor by how much the amount of annotated data for that class was increased from initial to final training data set. For simplicity, not all classes that the model was trained on are included.

| Behavior Type | TP Initial Model | TP Final Model | Label Data Increased |
| --- | --- | --- | --- |
| Ataxia | 0.79 | 0.83 | × 1.1 |
| Digging | 0.02 | 0.06 | × 4.3 |
| Eating | 0.38 | 0.88 | × 2 |
| Grooming | 0.17 | 0.79 | × 3.88 |
| Sniffing | 0.63 | 0.54 | × 5.64 |
| Startled | 0.08 | 0.09 | × 5.67 |
| Twitching | 0.03 | 0.29 | × 4.8 |
| Unsupported Rearing | 0.72 | 0.72 | × 1.7 |
| Catalepsy | - | 0.59 |  |
| Drinking | - | 0.91 |  |
| Gnawing | - | 0.29 |  |
| Interrupted Sleeping | - | 0.45 |  |
| Jumping | - | 0.85 |  |
| Supported Rearing | - | 0.70 |  |
| Stretched |  | 0.57 |  |
| Walking | - | 0.78 |  |
| Writhing | - | 0.11 |  |

## Conclusion

Our study has demonstrated that the integration of a human-in-the-loop approach, combined with advanced AI technologies, significantly enhances the efficiency and accuracy of rodent behavior analysis in the context of drug development.

The active learning component has proven instrumental in discovering and annotating rare behavior types, as evidenced by the substantial increase in annotations and the identification of several new behavior classes. This enhancement in data richness not only improves the model’s accuracy but also broadens the spectrum of behavior types that can be reliably detected. The increase in the amount of labeled data for rare behaviors has notably improved the model’s performance, as highlighted by the substantial improvements in TP rates for behaviors such as Grooming and Twitching. The system has been proven to accelerate the discovery process and aid labelers in prioritizing behaviors for annotation.

DeepRod represents a significant advancement in the field of automated rodent behavior analysis. Its ability to efficiently process large datasets, coupled with its enhanced detection and classification capabilities, makes it a powerful tool for drug discovery and development. The collected user feedback confirmed that the speed of annotating the experiment data and the discovery of such a large amount of new behavior types would not have been possible without the provided system. As the system continues to evolve, it holds great potential for further improving the understanding of rodent behavior, contributing to more effective and efficient drug development processes and safe clinical trials.

## Acknowledgements and contributions

Our deepest gratitude goes to Beatrice Kley, Cindy Janine Jung, and Michael Winter (Boehringer Ingelheim GmbH & Co. KG, Drug Discovery Sciences) for annotation of the data and for their feedback, wishes, ideas and immediate testing of these. We want to thank Malte Janssen, Moisei Shkil, Ziyad Sheebaelhamd, Lev Udaltsov, Atacan Korkmaz (Merantix Momentum GmbH) who developed software components used for DeepRod. DeepRod would not be possible without Florian Montel, Menorca Chaturvedi, and Markus Köster from opnMe.com [1] (F.M. Boehringer Ingelheim GmbH & Co. KG; M.C., M.K. Boehringer Ingelheim International GmbH). Fabian Heinemann and Peter Gross contributed the opnMe initiative at various stages and provided input on the requirements of the MVP. Kathrin Eschmann had a substantial impact on the late stage of the project and contributed to the manuscript (all Boehringer Ingelheim GmbH & Co. KG, Drug Discovery Sciences).

Adrian Loy, Miha Garafolj, Thomas Wollmann developed the machine learning method. Adrian Loy and Miha Garafolj did the algorithm implementation and performed the machine learning experiments. Hanna Behnke did the usability engineering and orchestrated the project. Cyrille Charnier and Philipp Schwarz evaluated the methodology. Georg Rast, Heike Schauerte contributed the initial concept and provided domain knowledge. Georg Rast and Thomas Wollmann supervised the project. Thomas Wollmann, Miha Garafolj, Adrian Loy, Heike Schauerte, Georg Rast, and Hanna Behnke wrote and proofread the manuscript.

## Ethical statement

Maintenance and handling of animals are carried out in compliance with (i) the ethical guidelines established by German National Animal Welfare Laws within the framework of the European Union Directive 2010/63/EU and (ii) the Guide for the Care and Use of Laboratory Animals produced by the National Research Council and the Association for Assessment and Accreditation of Laboratory Animal Care International (AAALAC). The study protocol was approved by the responsible German authority (Regierungspräsidium Tübingen).

